# Meniscus-enabled Projection Stereolithography (MAPS)

**DOI:** 10.1101/2023.06.12.544584

**Authors:** Puskal Kunwar, Arun Poudel, Ujjwal Aryal, Rui Xie, Zachary J Geffert, Haven Wittmann, Tsung Hsing Chiang, Mathew M. Maye, Zhen Li, Pranav Soman

## Abstract

Light-based additive manufacturing methods have been widely used to print high-resolution 3D structures for applications in tissue engineering, soft robotics, photonics, and microfluidics, among others. Despite this progress, multi-material printing with these methods remains challenging due to constraints associated with hardware modifications, control systems, cross-contaminations, waste, and resin properties. Here, we report a new printing platform coined Meniscus-enabled Projection Stereolithography (MAPS), a vat-free method that relies on generating and maintaining a resin meniscus between a crosslinked structure and bottom window and to print lateral, vertical, discrete, or gradient multi-material 3D structures with little-to-no cross-contamination or waste. We also show that MAPS is compatible with a wide range of resins and can print complex multi-material 3D structures without requiring specialized hardware, software, or complex washing protocols. MAPS’s ability to print structures with microscale variations in mechanical stiffness, opacity, surface energy, cell densities, and magnetic properties provides a generic method to make advanced materials for a broad range of applications.

## 1. Introduction

Current additive manufacturing methods are unable to match nature’s marvelous ability to arrange multiple materials in 3D configurations across scales to realize multifunctional structures.^[1–5]^ Extrusion or jetting methods with the use of multiple printing nozzles have been used to print multi-material structures, however, the print resolution, surface finish, shape fidelity, and speed are typically lower than vat photo-polymerization methods such as Projection stereolithography (PSLA) or digital light processing (DLP).^[6–8]^ A typical setup for PSLA consists of spatially modulated light patterns projected through a transparent bottom window to polymerize photosensitive liquid resin in a vat.^[9–11]^ The ever-growing library of resin formulations has already allowed PSLA to fabricate functional structures for applications in tissue engineering, soft robotics, photonics, and other fields.^[12–18]^ Despite this progress, multi-material printing with PSLA or similar methods remains challenging due to limits of hardware modifications, complex control systems, resin properties, and associated constraints as explained below.

Advances in PSLA technology such as Continuous liquid interface production (CLIP), high-area rapid printing (HARP), and Computed axial lithography (CAL), have improved printing speed, however, their ability to print multiple materials remain limited.^[19–21]^ Current multi-material PSLA methods continue to rely on proprietary hardware modifications such as the use of multiple vats, carousel-style rotators, linear movement of print stages, wiping mechanisms, and pressurized fluid flow to facilitate resin exchanges. ^[17,22–29]^ Instead of relying on passive refilling of resins, lateral (XY) stage motions in mask video projection-based stereolithography (MVP-SL)^[30]^ has been used to accelerate rapid refilling of resins, however this requires customized hardware and control modifications. Active perfusion of resin using viaducts embedded within the print geometry (Injection CLIP or iCLIP), has also been used to accelerate the refilling of resins.^[31,32]^ However, the embedded viaduct geometry itself induces artifacts to the printed construct. Multi-material printing with CAL, which rely on the projection of light patterns onto a rotating resin vat, is possible only when the second material is printed around pre-existing prints. Recently reported DLP-based centrifugal multi-material (CM) method involves lifting the stage out of the vat and applying centrifugal forces to clean resin residues from previous prints before re-immersing it back into the vat to print a second material; this requires substantial hardware modifications and is limited to discrete multi-material printing.^[33]^ Other strategies of greyscale light exposure or orthogonal dual-wavelength printing require customized resin photochemistry and/or complex optical engineering setups.^[34]^

Additionally, resin properties continue to play a dominant role in determining the printing capabilities of current methods. For instance, resins that are miscible with the lubricating liquid, necessary for rapid printing, cannot be used with HARP^[25]^. The DLP-based CM method is limited to materials that could withstand the centrifugal washing steps while CAL is limited to resins that exhibit high viscosity, high reactivity, and low scattering properties.^[33]^ Vat-free methods, that involve puddles of resins or smaller droplets with lateral or vertical stage movements, have also been used, although challenges related to separation forces between glass and polymerized parts, air-cleaning of high viscosity resins, and flow of large puddles remain^[35]^. In summary, despite recent advances, current multi-material PSLA methods require substantial hardware and software modifications, and key challenges related to resin refilling, contaminations during resin exchanges, and low recyclability due to unpredictable amounts of photoreactive components have not been addressed. These limitations become even more challenging for gradient printing, as generating consistent and accurate material transitions across the printed object become difficult. For instance, vat-based gradient printing suffers from poor control over the resin’s composition during the printing process, primarily due to issues like residue buildup and cross-contamination. These factors hinder the ability to precisely manipulate the resin’s properties required for successful gradient printing, emphasizing the need for new methods to overcome these challenges.^[36]^

In this work, we report the design and development of a Meniscus-enabled Projection Stereolithography (MAPS) platform. As compared to conventional methods, MAPS is a vat-free method that is capable of printing 3D structures with custom variations in lateral, vertical, discrete, or gradient properties with little-to-no cross-contamination or waste and does not require complex hardware or operating protocols. To demonstrate its potential for broad utility, MAPS was used to print 3D structures with custom variations in stiffness, opacity, surface energy, cell density, and magnetic properties.

## 2. Result and Discussion

### 2.1 MAPS

Conventional stereolithography methods rely on a resin reservoir or vat to additively crosslink layers with programmed stage movements. As compared to vat-based methods, MAPS relies on the generation and continuous maintenance of a resin meniscus between crosslinked layers and the bottom PDMS window. **(Figure 1A, S1)**. Initial formation of a meniscus occurs between liquid resin, bottom PDMS window, and air, thus creating a three-phase contact line modulated by tangential interfacial forces and surface tension. These combined forces result in the formation of a resin dome, with a contact angle denoted as θ1, between the resin and the PDMS window (**Figure 1A(i)**). As the print starts and progresses, another contact angle (θ2) is also formed between resin and crosslinked structures **(Figure 1A(ii))**. This leads to the formation of a resin meniscus (**Figure 1A(iii)**), which serves as a reservoir for the MAPS printing process.

**Figure 1.**
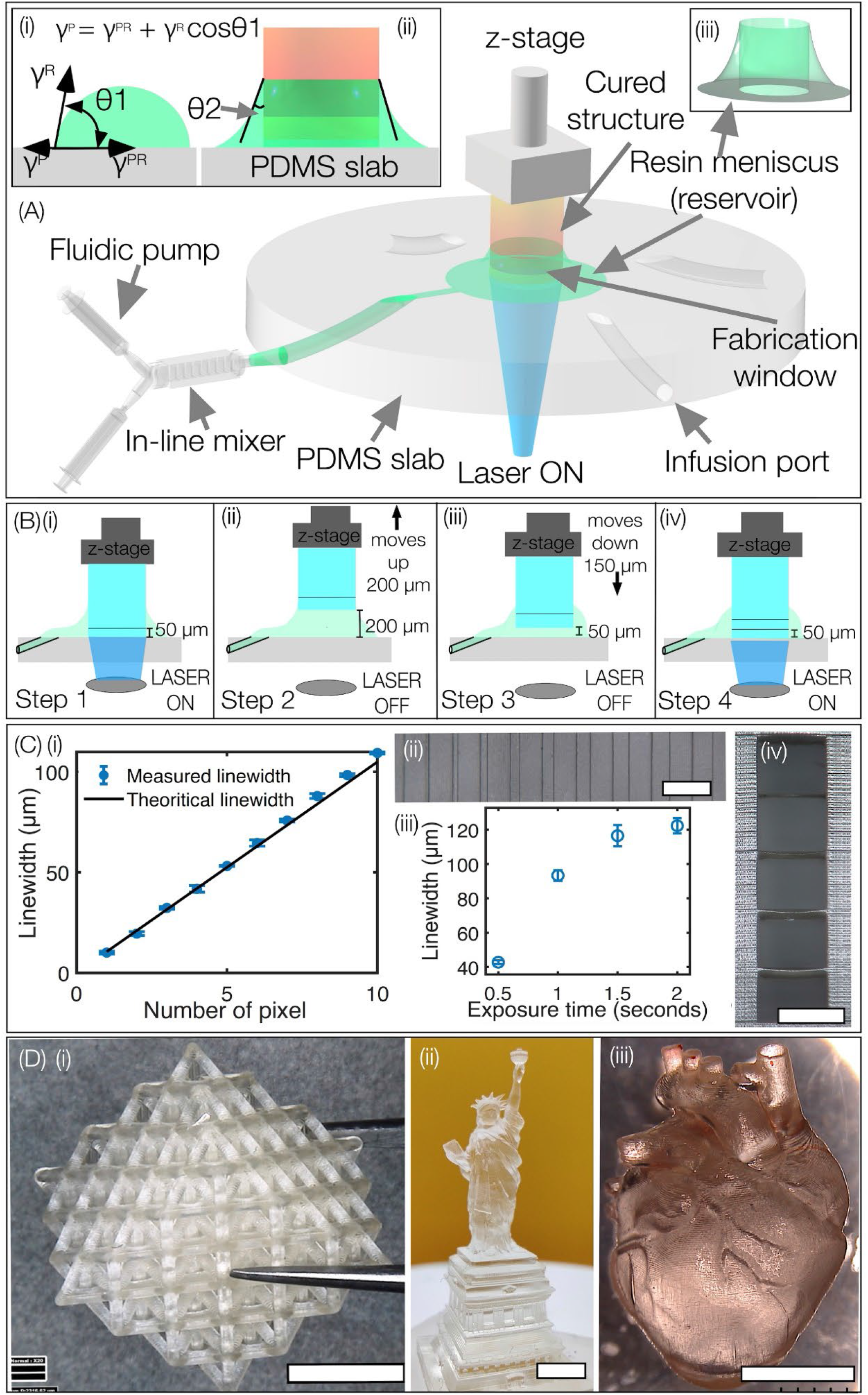
(A) Schematic setup illustrating the MAPS 3D printing by a continuous flow of resin solution using syringe pumps and an inline micromixer. (i) Formation of the three-phase contact line and resin dome due to resultant forces involved (γ^R^ - resin surface tension, γ^P^- PDMS surface tension, γ^PR^ - resin-PDMS boundary tension, and θ1 and θ2 = contact angles). (ii) Illustration of contact angle θ2 once the structure starts to print. (iii) Shape of the meniscus that acts as a reservoir for MAPS printing. (B)(i-iv) Process flow diagram of steps involved in MAPS. The up-down step elevation of the z-stage draws the liquid into the fabrication area, while the projection of spatially patterned 405 nm light crosslinks the resin solution. (C) Plots and photographs showing the (i,ii) lateral (scale bar-200 µm) and (iii, iv)axial resolutions (scale bar - 500 µm) of the printed structure using PEGDA 400 MW. (D) (i) Lattice cube (scale bar-3 mm) (ii) the Statue of Liberty (scale bar - 3 mm) and (iii) human heart structure (scale bar - 3 mm) are fabricated using MAPS.

A 405 nm light, spatially modulated via a digital micromirror device (DMD) and projected through the PDMS bottom window, is used to crosslink liquid resin in a layer-by-layer manner (**Figure S1**). Vertical stage movement generates a suction force that continuously draws the resin from the meniscus to the fabrication window (light projected area on the PDMS window). As the printing process continues, a fluidic pump is used to replenish the resin to ensure the maintenance of the resin meniscus throughout the printing process (**Figure 1A**). Most of the suction force is utilized to pull the resin against the PDMS interface, while a fraction of force is also applied against the adhesion of the interface between the resin solution and the crosslinked structure. The infusion flow rate of new liquid resin is matched with the suction force to ensure continued maintenance of the meniscus throughout the printing process. Modular use of multiple infusion ports, syringe pumps, and inline micromixers, allow easy and rapid configuration of the system for printing customized multi-material structures.

In MAPS, similar to other PSLA methods, the oxygen permeability of the PDMS window facilitates the generation of a ‘dead zone’ just above the fabrication window to prevent unwanted adhesion of newly crosslinked layers to the bottom window during the printing process. (**Figure S1A)**. For a particular resin formulation and CAD design, the upward stage movement was coordinated with layer exposure times, dead-zone thickness, light dosage per layer, and the fluidic flow rate of new resin. Before multi-material printing, model resin PEGDA 400 MW was used to quantify lateral and axial resolution of MAPS using a ‘line array’ and ‘staircase’ templates respectively. The desired layer thickness (50 µm, for PEGDA 400) was generated by programming the stage to move up (200 µm) and down (150 µm) during light irradiation. (**Figure 1B(i-iv)**). Lateral and axial resolutions were quantified as 10.05 ± 0.7 µm and 42.79 ± 1.12 µm respectively. For vertical resolution experiments, the PEGDA resin was mixed with 1% ITX (photoabsorber) and 0.25% Irgacure (photoinitiator) to achieve a range of axial feature size ranging from 42.79 ± 1.12 to 122.24 ± 4.45 µm using exposure times varying from 0.5-2 seconds per layer (**Figure 1C(i-iv)**). This formulation was used to print a ‘lattice cube’, ‘Statue of Liberty’, and ‘human heart’ structures using a layer thickness of 50 µm, a laser intensity of 3.25 mW/cm^2^ and exposure time of 0.7 seconds per layer. The printed structure was washed and developed in an ethanol solution for 5 minutes. (**Figure 1D(i-iii), Video V1**).

Several commercially available resins such as Black resin, Flexible-X, Photocentric grey and Composite-X were also tested **(Figure S3)**. Details related to resin formulation and associated printing conditions can be found in **Table S1**. Overall, these results demonstrate the capability of MAPS to effectively print 3D structures with resolutions at par with other vat-based methods.

### 2.2 Optimization of MAPS for multi-material and gradient printing

Maintaining meticulous control over the flow rate and small meniscus volume is critical for achieving successful multi-material discrete and gradient 3D printing where resin properties continuously vary during the printing process. (**Figure 2A)**. To ensure an uninterrupted printing process, it is important to monitor and tune the infusion flow required to overcome the resistance offered by the PDMS surface. (**Figure 2A(i,ii)**). To avoid cross-contamination during resin exchanges, a minimum volume of resin should be present in the meniscus; higher volumes lead to undesired mixing of resins or accumulation of resin around the crosslinked layers while low volumes lead to interrupted printing and associated defects. To address this challenge, we used a multiphase many-body dissipative particle dynamics (mDPD) model to simulate the dynamic process of MAPS **(Figure 2B, Video V2)**. The setup shows the fabrication window (between z-stage printhead size, D = 5 mm), PDMS window, and infusion port (Dt = 0.5 mm). Results for different substrate wetting contact angles (θ1 = 20°, 45°, 90°, and 110°) showed that when the substrate becomes more hydrophilic, liquid adhesion is increased, resulting in the spreading of the droplet onto a wider wetting area. On the other hand, increased hydrophobicity of the substrate (θ1=110°) weakens liquid adhesion and makes it more susceptible to liquid-bridge breakage at a location between the printhead and pump outlet. Multiphase fluid simulations provided valuable insight into the underlying physics of multiphase flow and droplet wetting. By visualizing and analyzing the multiphase flow dynamics, we identified an ideal contact angle range (30°-45°) for multi-material printing without flow disruptions/breakage and ensuring a small volume of resin at the meniscus.

**Figure 2.**
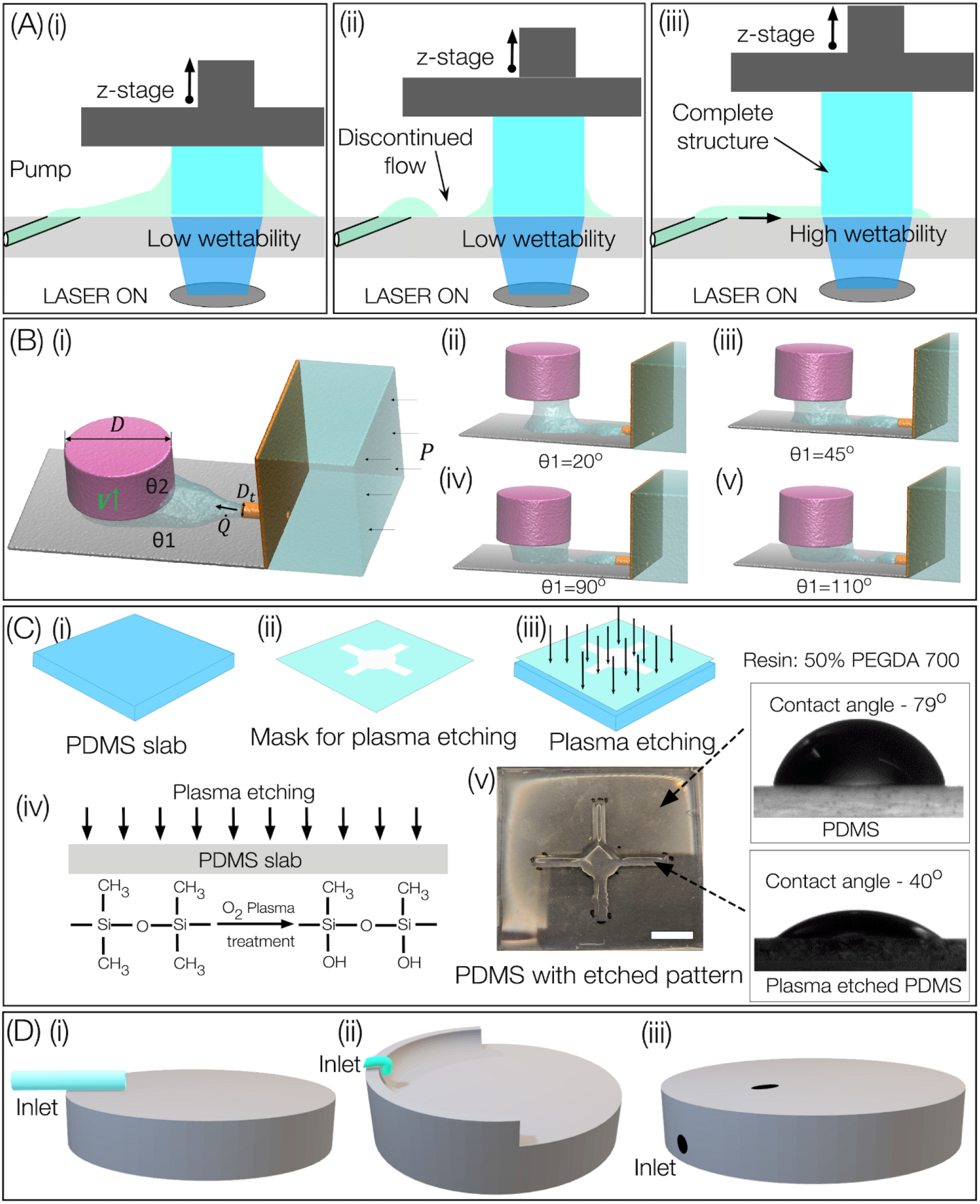
(A) Schematic diagram of (i) flow of resin to the fabrication area with low wettability (ii) breakage of flow during the printing process (iii) continuous flow of resin to the fabrication area on surface with high wettability. (B)(i) System setup of the computational fluid model for MAPS setup with PDMS substrate with different wetting contact angles. Simulation results show high substrate wettability is better suited for the uninterrupted MAPS process. (C) Patterning hydrophilicity on PDMS window using oxygen plasma treatment. Change in contact angles for 50% PEGDA 700MW resin is depicted (Scale bar - 5 mm). (D) Different configurations of sample holders were used in this study. (i) Side tube configuration. (ii) Top tube configuration. (iii) Side channel configuration with the slanting angle design.

For the resins we used, typical contact angles on PDMS window were in the range of 55°-90°. To achieve the ideal contact angle range, high wettability areas were patterned onto the PDMS window to enable easy flow of resin from the infusion port to the fabrication area without any disruptions. The high wettability area was patterned by plasma etching (5 minutes; RF power-75W) PDMS window with a cross-shaped mask. **(Figure 2C(i-iv))**. The high wettability pattern facilitates low-volume resin flow from infusion ports toward the fabrication area. The contact angle of patterned areas was in the range of 37°-55° for all the polymers tested (**Figure S4, S5**). Plasma-etched flow paths facilitated drawing resins from greater distances and enable multi-material and gradient printing with minimal cross-contamination and waste. Three distinct configurations of PDMS window and infusion ports were tested. (**Figure 2D(i-ii)**. While all designs worked with MAPS, the third, easy-to-clean, stage design was primarily used in this work.

### 2.3. Multi-material printing using MAPS

Here, we demonstrated the versatility of MAPS printing by designing and printing a variety of multi-material structures using two independent infusion ports with programmable flow rates. The flat multi-port configuration was used to enable easy resin exchanges with low or no cross-contaminations in both vertical and lateral directions **(Figure 3A)**. For instance, ‘The Statue of Liberty’ (**Figure 3B(i), Video V3)**, and ‘a knotted vessel’ geometries were printed with discrete sections of PEGDA 250 MW and Photocentric grey resin in the vertical directions (**Figure 3B(ii))**. Rapid cleaning steps were introduced between resin exchanges, and ‘Statue of Liberty’ and ‘lattice cuboid’ geometries were printed using PEGDA 250 MW (transparent) and commercially sourced Black resin (black, opaque). Results show distinct layers without any contamination between layers **(Figure 3C(i,ii))**. Simultaneous lateral and vertical multi-material printing was demonstrated by printing an array of 3D pillars with a checkerboard roof pattern using Photocentric grey resin and Composite-X resin (**Figure 3C(iii)**). A monolithic log-pile structure was printed with discrete sections of PEGDA 700 (top) and PEGDA 6k (Bottom) (**Figure 3C(iv))**. PEGDA 700 (stiff, top) section retained its shape with distinct edges, while PEGDA 6k (soft, bottom) section swelled when submerged in DI water and collapsed upon drying in air. Cell-adhesive Gelatin Methacrylate (GelMA) in the form of ‘Syracuse University’ SU logo was printed on top of inert synthetic PEGDA 6k hydrogel using MAPS. Human osteosarcoma cells (Saos-2) were seeded on the construct and labeled for F-actin to visualize cell morphology. Results show high adhesion and spreading of cells only on GelMA regions (SU logo) while minimal-to-no cell attachment was observed on inert PEGDA 6k MW slab. **(Figure 3C(v))**. MAPS was also used to print a 3D structure with alternating hydrophobic (PEGDA 250) and hydrophilic (PEGDA 6k) patterns forcing a water drop to travel out-of-focus along the hydrophilic line pattern, as compared to control samples **(Video V4)**. Lastly, a three-chambered microfluidic device separated by micro-post arrays was printed with a central ‘hydrophilic’ (PEGDA 6k MW) channel flanked by two ‘hydrophobic’ (PEGDA 250 MW) channels using MAPS. Results show that green dye mixed in DI water remains within the central ‘hydrophilic’ channel and does not leak into the ‘hydrophobic’ channels. In contrast, an identical device printed only with hydrophobic resin results in leakage of dye in all channels. Control over leakage in such devices can be useful for organ-on-a-chip applications.

**Figure 3.**
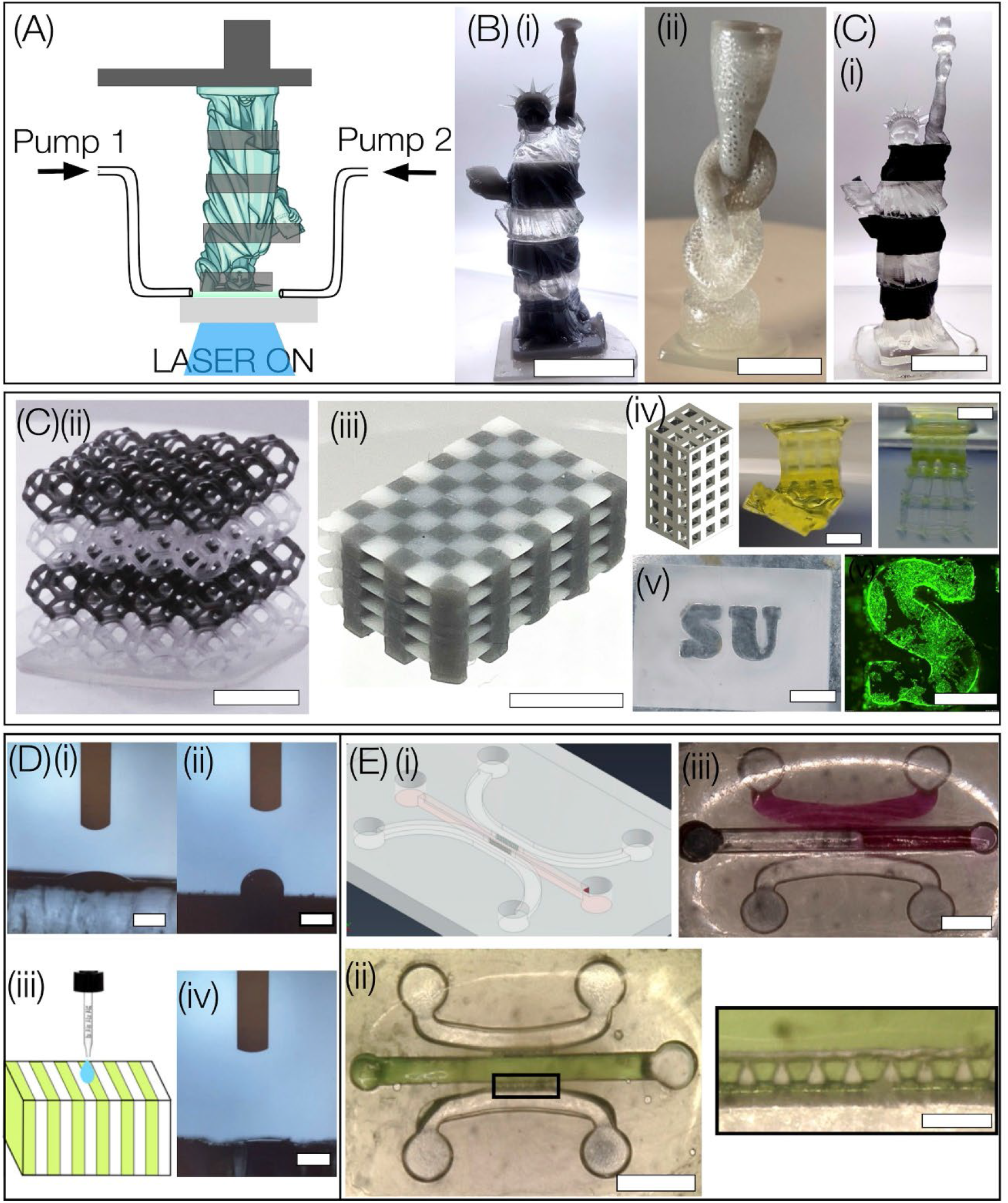
Multi-material printing using MAPS. (A) Configuration with two flow pumps. (B) Multimaterial printing without cleaning steps: (i) Statue of Liberty (scale bar - 5 mm) and (ii) knotted vessel printed using PEGDA 250 and Photocentric grey resin (scale bar - 5 mm), and (C) with cleaning steps between resin exchanges (i) Statue of Liberty printed using PEGDA 250 MW and Black resin (scale bar - 5 mm). (ii, iii) Lattice cuboid and arrays of 3D pillars with checkerboard roof (scale bar - 5 mm). (iv) Multi-material structure with hard (90% PEGDA 700) and soft regions (10% PEGDA 6k); only soft regions swell upon hydration and collapse when exposed to air. (Scale bar - 2 mm). (v) Multi-material printing of biologically inert (10% PEGDA 6k) and cell adhesive material (10% GelMA) (Scale bar - 2 mm), with attachment and proliferation of cells only on GelMA (Scale bar - 1 mm). (D) (i, ii) Contact angle formed by water drop on flat slabs printed with PEGDA 6k, and PEGDA 250. (iii) Schematic and (iv) printed structure composed of alternating layers of hydrophobic (PEGDA 250) and hydrophilic (PEGDA 6k), and associated contact angle (Scale bar - 1 mm). (E)(i) CAD design and (ii) 3D printed three-channel hydrophobic (PEGDA 250) microfluidic device with hydrophilic (PEGDA 6k) central channel separated by an array of micro-posts (scale bar – 3 mm). A zoomed-in section shown by a black rectangle depicts an array of micro-post (scale – 500 µm). Green dye perfused into the central channel does not leak out into the side channels, as compared to red dye perfused into (iii) an identical chip printed using single material (PEGDA 250) (Scale bar - 3 mm).

### 2.4. Gradient printing using MAPS

To extend the capability of MAPS to print 3D structures with customized gradient properties, an inline micromixer was inserted between two syringe pumps and one infusion port of PDMS window. (**Figure 1A, S6A)**. Details related to the design, fabrication, and performance characterization of the micromixer are provided in the SI (**Figure S6, S7, Video V5**). Based on the desired gradient properties, multiple resins were mixed at defined flow rates and concentrations to generate droplets within the fabrication window. During the MAPS process, laser exposure time per layer was modulated based on mixed resin formulation. The capabilities of printing gradient complex 3D structures were assessed using a range of structures, materials, and configurations. For instance, ‘twisted tower’ geometry with a defined color gradient (top-green, and bottom-yellow) was printed using PEGDA 700 resin **(Figure 4A(i-ii))**. A combined mechanical and color gradient ‘Twisted tower’ structure was also printed using PEGDA 700 (brown, stiff) and PEGDA 6k (colorless, soft) with flow rates varying between 0-0.2 ml/min for both resins in the opposite direction and at the same time the exposure time from 0.8-2 seconds to obtain identical curing depth per layer despite changes in the mixed resin droplet formulations (**Figure 4B(i), Video V6**). Next, the same structure was printed by reversing the flow rate of the two resins to generate mechanical and color gradients in opposite directions (**Figure 4B(ii))**. Flowrate of the resins and associated exposure time are also shown (**Figure 4B(iii))**. To assess the mechanical properties of the gradient structure, different sections (∼400 µm thick) marked by black boxes in **Figure 4B(ii, iv)**, were used for standard compression tests. Results show that the modulus varied from 0.249 (bottom) to 3.9 MPa (top) along the height of the structure. **Video V7** shows this top-heavy gradient structure vibrates even upon gentle perturbation. To demonstrate the use of MAPS with commercially available resins, we printed gradient ‘Statue of Liberty’ structures using a combination of PEGDA 250 (colorless) and photo-centric grey DLP resin by linearly varying flow rates (0-0.2 ml/min). Both front and back views of the printed structure (along with associated flow rate and exposure time) are shown to highlight the uniform distribution of the resin composition achieved with this method (**Figure 4C(i-iii))**. We also printed the same structure with exponentially varying flow rates (**Figure 4D)**. The modularity of choosing multiple ports, micromixers, resin flow rates, and concentrations underscores the unique capabilities of MAPS to print 3D geometries with customized gradient properties.

**Figure 4.**
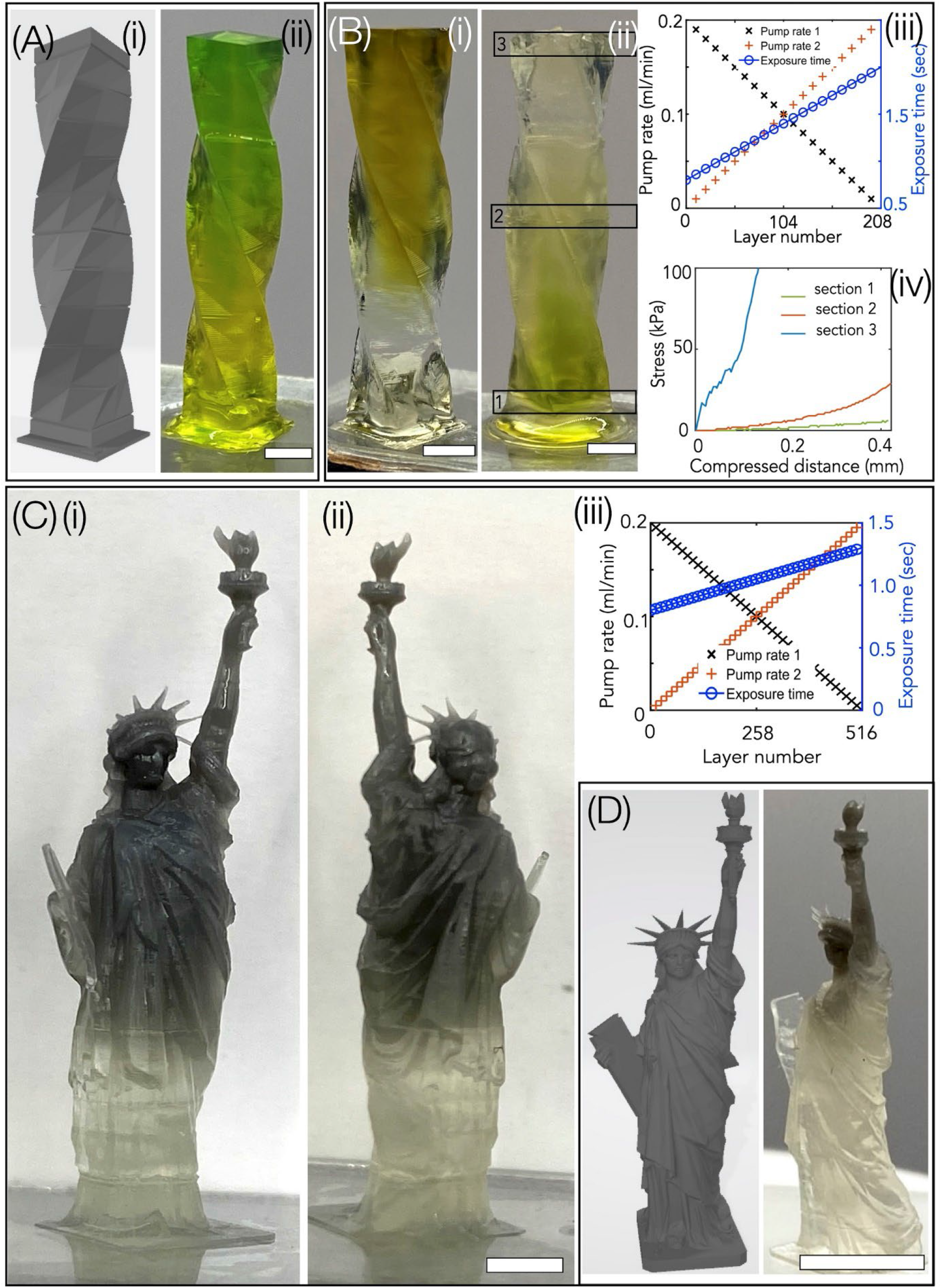
Gradient property printing with MAPS. (A) (i) displays the CAD design of the twisted tower. (ii) exhibits the two-color gradient printing of the twisted tower using green and yellow color PEGDA 700 hydrogels (scale bar - 2 mm). (B) (i-iii) exhibits the twisted tower with dual color and stiffness gradients with PEGDA 700 and PEGDA 6k resins with associated changes in infusion flow rates and exposure times. (Scale bar - 2 mm). (iv) Stress-strain plots from different sections of the printed structure (black rectangle) where modulus varied from 0.249 (bottom) to 3.9 MPa (top). (C) (i-iii) displays the front and back views of the gradient liberty tower structure printed using PEGDA 250 and photo-centric grey with associated flow rates and exposure times (scale bar - 5 mm). (D) shows a CAD design of the structure (Statue of Liberty) and multi-material gradient structure with exponentially varying flow rate (Scale bar - 10 mm).

### 2.5. Gradient printing with additives (magnetic nanoparticles, living cells, carbon black)

Here, we demonstrated MAPS’s capability to print 3D gradient structures with a range of additives **(Figure 5)**. First, iron oxide magnetic nanoparticles (NPs) were chosen as the model NP due to its broad use in medicine, biosensing, catalysis, agriculture, and the environment. ^[37,38]^ Before performing gradient printing with NPs, processing conditions were optimized for mixed resin formulations. For instance, MAPS printing with a light intensity of 3.25 mW/cm^2^ resulted in a curing depth of 120 µm for resin with 10% NPs and an exposure time per layer of 25 sec, as compared to a curing depth of 150 µm and an exposure time of 1.5 sec for only resin. Optimized parameters were used to print a cylindrical geometry with a gradient structure with a maximum concentration of 10% NPs in the top section and decreasing NP concentration along the length of the cylinder. Results show that only the structures with NPs respond to the magnetic field, and the response of the gradient-aligned NPs structure was distinct from that of the structure with uniform NP distribution **(Figure 5A(i-iv))**. The ability to incorporate other functional NPs in customized gradient configurations can be broadly applied to many applications ranging from structural colors, soft robotics, and photonics to name a few.

**Figure 5.**
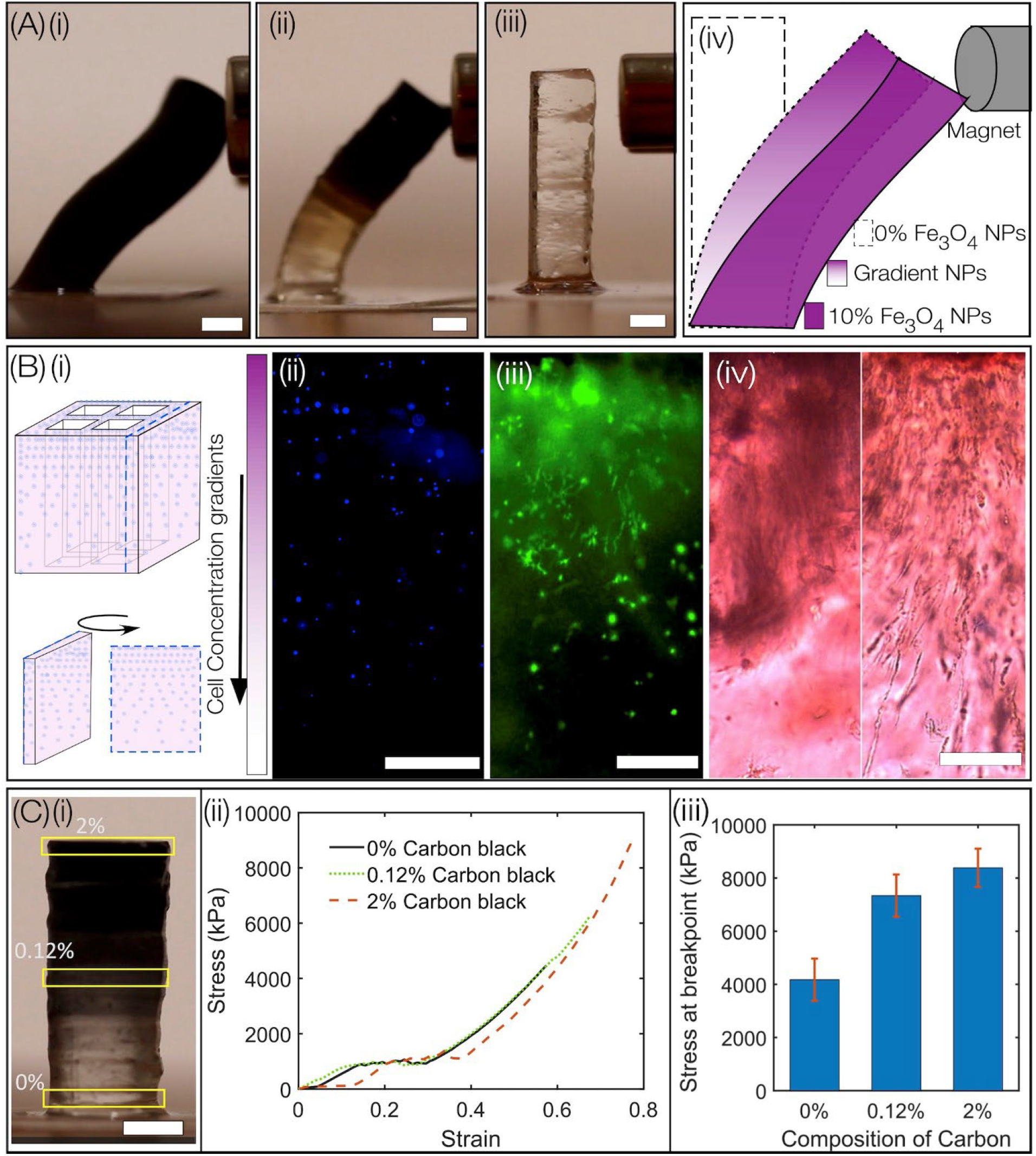
A(i-iv) Response of cylinder geometry with uniformly distributed magnetic NPs, gradient NPs, and no NPs to the magnetic field. (Scale bars-2 mm). (B)(i) Schematic representation of the bioprinted geometry using MAPS depicting gradient cell densities along the z-direction. Post-printing, (ii) the DAPI-stained construct cultured for 3 days was sliced, and a section was viewed under a fluorescence microscope. (iii) F-actin staining of the sliced construct (iv) Demonstration of the gradient distribution of bone mineral in the sliced construct cultured for 14 days in osteogenic media (Scale bar (ii-iv) -500 µm). (C)(i) Cylindrical structure printed using a resin with Carbon black NPs additive (Scale bar-2 mm). (ii-iii) Compressive stress-strain plot of the cylinder section marked by yellow rectangles, and associated stresses.

Second, MAPS utility for bioprinting application is demonstrated. Although a range of bioprinting methods exist, printing structures using gradient cell densities remains challenging.^[36,39–41]^ Here, MAPS was used to print a 3D structure with a gradient of cell density using a bioink composed of 5% GelMA, 4% PEGDA 6k, and 0.25% LAP, and model cells (MC3T3 pre-osteoblasts, 1M/ml). Optimized conditions (laser power: 3.6 mW/cm^2^; exposure time: 1 seconds/layer; layer thickness: 50 µm) were used to bioprint a 3D structure with gradient densities of MC3T3 along the z-direction (from top to bottom) **(Figure 5B)**. Post-printing, the sample was washed with PBS three times, sliced in half, and stained for nuclei (DAPI) and F-actin, which revealed a gradient of cells with high spreading on the top section and almost no cells on the bottom. (**Figure 5B(iii)**). The printed sample was cultured under osteogenic media for 14 days and stained with Alizarin Red to identify regions of mineral formation. Results show a gradient of mineralization from the top to the bottom section of the printed sample. (**Figure 5B(iv))**. MAPS bioprinting can be extended to tissue-specific gradients as well as multi-cellular gradient constructs for potential tissue engineering and regenerative medicine applications.^[42,43]^

Third, MAPS gradient printing was demonstrated using ‘carbon black’ as the additive. Carbon black as an additive reinforces the mechanical properties of the structure.^[44,45]^ A cylindrical structure is printed using a PEGDA 700 with carbon black, where the concentration of carbon black varies from 0% to 2% (**Figure 5C(i))**. Sections of the structure (marked by yellow rectangles) were compressed to obtain a compressive stress-strain plot, which provides a clear indication of the gradient and enhanced mechanical properties of the printed structure (**Figure 5C(ii-iii)**). Results show gradient changes in breakpoint stress, from 8385 ± 719 kPa to 4174 ± 793 kPa for 2 to 0 wt% carbon black.

## 3. Conclusion

MAPS, a new vat-free method that relies on the generation and maintenance of a small amount of resin meniscus in the fabrication window, overcome many challenges related to current light-based multi-material printing. Here, we show that MAPS can print lateral, vertical, discrete, or gradient multi-material 3D structures using a wide range of resins with minimal cross-contamination or waste and without the use of specialized hardware, software, or complex materials exchange and washing protocols. MAPS’ ability to print multi-material 3D structures, with custom variations in mechanical stiffness, opacity, surface energy, cell densities, and magnetic properties, opens new possibilities to make advanced materials with customized properties.

## 4. Experimental Section /Methods

A detailed method section is provided either within the manuscript or in the supplementary information (SI).

## Supporting Information

Supporting Information is available from the Wiley Online Library or from the author.

## Acknowledgment

The authors would like to express their gratitude to Daniel A. Fougnier for his valuable contribution in measuring the viscosity of various photo resins. Additionally, the authors would like to extend their thanks to Ashraf Tariq Alnatour for his assistance in programming the pump. Financial support for this project was provided by the National Institutes of Health (R21 GM141573-01) and the Syracuse University Collaboration of Unprecedented Success and Excellence (CUSE) and the BioInspired Seed grant program.

Received: ((will be filled in by the editorial staff))

Revised: ((will be filled in by the editorial staff))

Published online: ((will be filled in by the editorial staff))

