## Supplementary data for "Meniscus-enabled Projection Stereolithography (MAPS)"

### S1. Synthesis of resins

This study employed various resin formulations as described below:

**(1) Gelatin Methacrylate (GelMA)** prepolymer solutions with 7.5% w/w were prepared by dissolving the required dry macromer mass in Millipore water with 1.0% (w/w) LAP (photo-initiator). GelMA synthesis was performed using a previously reported protocol.[1] In brief, 10 gm of porcine skin gelatin (Sigma Aldrich, St. Louis, MO) was mixed in 200 ml of phosphate-buffered saline (PBS, Thermo Fisher Scientific) at 45°C, and methacrylic anhydride was added to the solution, which was stirred for 3 hours. The resulting mixture was then dialyzed against distilled water for 1 week at 40°C to remove any unreacted groups. The dialyzed GelMA was lyophilized in a freeze dryer (Labconco, Kansas City, MO) for one week. To prepare a 15% (w/v) GelMA stock solution, 1.5 g of freeze-dried GelMA was mixed with 10 ml of deionized water (dissolved at 40°C), and 0.25% (w/v) UV photoinitiator lithium phenyl-2,4,6-trimethyl-benzoyl phosphinate (LAP) was added. The GelMA pre-polymer solution was then diluted using DI water to obtain either 7% or 10% GelMA, filtered (pore size=0.2 μm), and used within 2 hours after preparation.

**(2) LAP synthesis.** Lithium phenyl-2,4,6-trimethyl-benzoyl phosphinate (LAP) was synthesized in two steps according to a published procedure.[2] First, under argon at room temperature, 2,4,6-trimethyl benzoyl chloride (4.5 gm, 25 mmol) was added dropwise to dimethyl phenyl phosphonate (4.2 gm, 25 mmol) with continuous stirring. The reaction mixture was stirred for 24 hours, after which an excess of lithium bromide (2.4 gm, 28 mmol) in 50 mL of 2-butanone was added to the reaction mixture. The mixture was then heated to 50°C and allowed to stir for 10 minutes, resulting in the formation of a solid precipitate. After cooling the mixture to room temperature and letting it rest overnight, it was filtered, and the filtrate was washed with 2-butanone (3 x 25 mL) to remove any unreacted lithium bromide. The resulting product was then dried under vacuum to yield LAP (6.2 g, 22 mmol, 88%) as a white solid.

**(3) PEGDA (6000 MW*)*** was synthesized in the laboratory and dissolved in water to achieve 10% w/w PEGDA 6k, with 1% LAP w/w.[3] To synthesize PEGDA 6k, the first step involved preparing anhydrous dichloromethane (DCM) by refluxing it over calcium hydride for 2 hours, followed by simple distillation and storage over activated 3A molecular sieves. Next, 18 gm of PEG 6k was dissolved in 300 ml of anhydrous DCM and cooled in an ice bath. Under a nitrogen blanket with stirring, 1.69 ml of triethylamine was added to the chilled solution, followed by the dropwise addition of a solution of 0.975 ml acryloyl chloride diluted to a final volume of 15 ml in anhydrous DCM via an addition funnel. The reaction mixture was allowed to stir for 30 minutes under a nitrogen blanket, and then allowed to proceed overnight with constant stirring in a covered flask under nitrogen. Afterward, the reaction mixture was vacuum filtered through Celite, followed by washing of the Celite with three aliquots of DCM. The excess DCM was removed using a rotary evaporator until the concentrate became slightly cloudy. The concentrate was reconstituted with a minimal amount of fresh DCM to achieve a clear solution. PEGDA was then precipitated by dropwise addition to hexane with vigorous stirring. The resulting precipitate was isolated via vacuum filtration, washed with three aliquots of hexane, and then three aliquots of -80˚C diethylether. Finally, the washed PEGDA was dried under a vacuum at room temperature for two days.

**(4) PEGDA (700 MW;** Sigma Aldrich) solution was formed by dissolving 100% solid macromer in its waxy form in millipore water, resulting in 20% w/w with 0.2% LAP (w/w).

For all (1-4) resin formulations, Tartrazine was used as a photo absorber in a concentration of 10% of LAP (w/w). The solutions were usually vortexed and degassed overnight at 37ºC before use.

**(5) PEGDA (250 MW**; Sigma Aldrich) was added with 0.5% ITX and 0.25% Irgacure w/w and used without dilution. **(6) PEGDA solution (MW** 400; Sigma Aldrich) was prepared by adding the purchased product to a final concentration of 100%, with 0.5% ITX and 0.25% Irgacure. PEGDA solutions were vortexed and degassed before use.

**(7) Commercial resins.** A Black resin (B9 creations), Flexible-X resin and Composit-X resin (Liqcreate), and photocentric grey resin (MatterHackers), were used without further modification.

### S2. Surface modification of glass substrate

To ensure adhesion of a crosslinked structure to the glass, glass coverslips (18 mm x 18 mm, No.2; Globe Scientific) were surface modified with a methacrylate function group. A cleaned coverslip was first treated in a Piranha solution (a mixture of sulfuric acid and hydrogen peroxide in a 7:3 ratio) for 20 minutes. The glass was then washed with MilliPore water and 100% ethanol (Fisher Scientific, Pittsburgh, PA), and dried using compressed air. To functionalize the coverslip, it was dipped in a bath containing 85 mM of 3-(trimethoxysilyl) propyl methacrylate (Fluka, St. Louis, MO) in ethanol with acetic acid (pH 4.5) overnight at room temperature. Finally, the coverslips were washed in 100% ethanol and dried. This process ensured that the coverslips had a methacrylate function group on the surface, allowing for strong adhesion of the crosslinked structure to the glass.

### S3. Optical Setup

As compared to conventional vat-based PSLA methods (**Figure S1 A(i)**), vat-free MAPS is schematically depicted in **Figure S1 A(ii)**, **B**. This approach employed a layer-by-layer additive mode of fabrication, utilizing a 405 nm laser source (Toptica) that can generate a continuous-wave (CW) laser beam with a maximum power output of 300 mW. To collimate and expand the laser beam, a shutter (SH05, Thorlabs) was positioned after the laser source, and a 2f-transfer lens assembly (f=40 mm and 200 mm) is utilized. The laser beam was spatially filtered through a 25 μm pinhole, which was then directed through a diffuser (RPC Photonics Inc) to transform the Gaussian intensity distribution of the laser beam to a uniform distribution. The diffuser was mounted on a rotating mount to minimize the laser speckle generated by the diffuser. The laser beam was collimated using a convex lens and directed toward a Digital Micromirror Device (DMD) (0.95" 1080p UV DMD, DLi Inc), which is an array of micromirrors that can be modulated to spatially pattern laser beams. The patterned laser beam was projected onto infinity-corrected projection optics, which include two lens systems (f=300 mm and f=300 mm at a distance of 60 cm). A precisely adjusted distance between the lenses focused the spatially patterned within the fabrication window. The PDMS slab, was positioned above a heater (WP-16 Warner instrument) that had a hole in the center (not shown in the figure). The heater was used to warm the PDMS membrane to 40ºC for only GelMA printing, as GelMA typically gels at room temperature. The L-shaped stage and its movement were controlled using a three-dimensional linear stage (L505, PI). A custom LabVIEW code was used to coordinate the alteration of the mask in the DMD with the movement of the stage in the z-direction. Additionally, two syringe pumps from New Era Pump Systems Inc. and a custom-designed microfluidic mixer were employed to provide a continuous supply of resin. The SyringepumpproV1 software was utilized to program the pumps during the printing of multiple materials and gradients structures.

**Process Flow.** The process of creating a 3D structure involves several steps. First, a CAD model was designed in Illustrator or Solid Design software. Next, MATLAB code was used to slice the 3D structure into 2D png images to generate a digital mask that corresponds to individual printing layers. Each image was then processed with a MATLAB code to create a 1-bit image with the desired spatial distribution for its unique 2D slice. These images were fed into a custom-written LabVIEW code that synchronized the pump flow rate, volume, upward movement of the stage, and exposure of the light pattern generated by DMD. This process ensured precise control over the printing of each layer of the 3D structure. The CAD design and dimensions of the single-channel and four-channel sample holder with side channel configuration are depicted **(Figure S2(A-C)).**

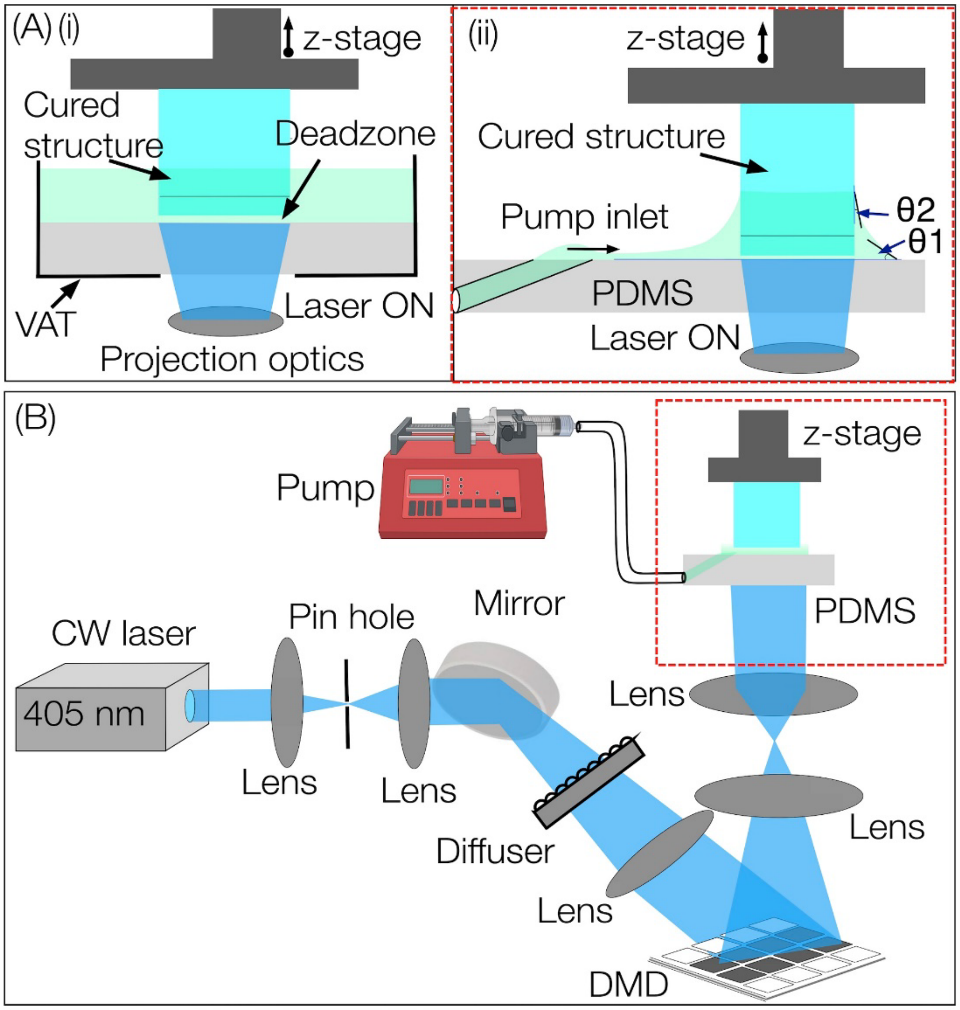


**Figure S1. (A)** (i) Illustration of conventional vat-based PSLA with bottoms-up configuration. Here spatially modulated light patterns are projected through a transparent bottom window to polymerize photosensitive liquid resin in a vat in discrete layers or in a continuous fashion to print the final 3D structure. (ii) Schematic illustrating the vat-less MAPS printing of 3D structure by a continuous flow of resin droplets using programmable syringe pumps. The figure depicts the meniscus governed by two contact angles (θ1 and θ2). θ1 is the contact angle formed by resin on PDMS, while θ2 is formed by resin on the crosslinked structure. **(B)** The custom-built MAPS printing setup consists of a 405 nm laser system, 2f lens-telescope system, rotating diffuser for converting the Gaussian intensity distribution of light to hat-shaped intensity, Digital Micromirror Device (DMD), projection optics, polymer holder with PDMS as bottom window, and stage mounted on programmed z-stage which is translated in the vertically upward direction during the printing process.

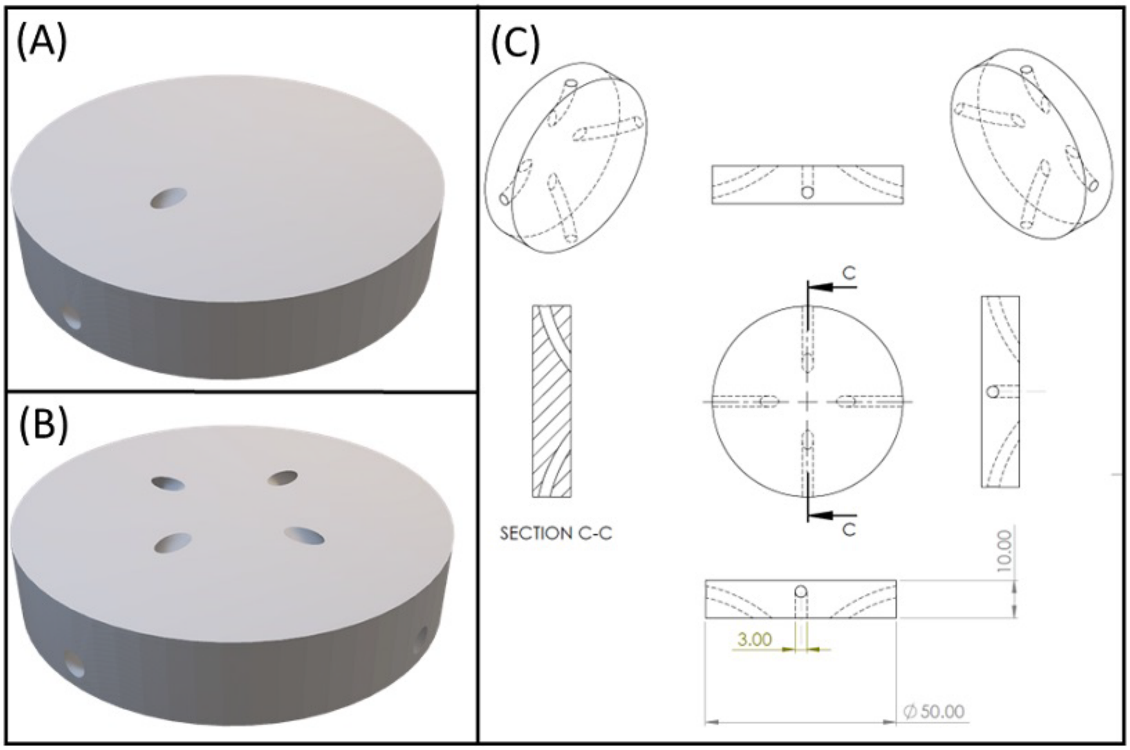


**Figure S2**.CAD design of (A) single-channel and (B) four-channel sample holder. (C) Detailed dimensional and design of four-channel sample holder

**S4. Characterization of XYZ resolution of MAPS using model resin**

In this study, we investigated the lateral (xy) and axial (z) resolutions of the MAPS system using PEGDA 400 resin. These resolutions depend on several factors, including the printing system's light exposure conditions, z-direction displacement of the stage, and material properties such as transparency, monomer reactivity, radical diffusion, and photochemical efficiency. PEGDA 400 photopolymer is proven to provide high-resolution 3D printed structures. To determine the lateral resolution, we printed lines of varying widths (1-10 pixels) from a digital mask. The resulting array of lines with linewidths ranging from 10.05 ± 0.7 µm to 109.42 ± 0.54 µm is depicted (**Figure 1C (i, ii)**). The smallest feature size (10 µm) was achieved using a 1-pixel line, which closely corresponds to the theoretical resolution determined by an individual mirror's size of DMD. We plotted the theoretical feature size against the experimental linewidth of the structures printed with a laser intensity of 3.25 $mW/cm^2$ and exposure time of 1 second, which showed good agreement between theory and experiment. The axial resolution, or z-direction resolution, depends largely on the curing depth, which is the thickness of the cross-linked layer in relation to the irradiation dose. The curing depth is affected by the z-direction motion of the stage, resins' optical absorbance, and crosslinking kinetics. Excessive light penetration beyond the desired curing depth results in unwanted crosslinking, leading to artifacts, particularly while printing channels, undercuts, and overcuts. To address this issue, we aimed to decrease the curing depth by increasing the photopolymer's absorbance using various methods, such as adjusting light dosage and using photo absorbers like Tartrazine, Sudan, and TINUVIN. In this study, we used ITX as a photo absorber, which also acts as a photosensitizer. However, ITX's photosensitizer efficiency in acrylate-based resin is low, so the primary role of ITX in our study was to limit the light's penetration depth to achieve the desired curing depth. We printed a ladder structure with varying exposure times from 0.5 to 2 seconds while maintaining a constant exposure intensity of 3.25 $mW/cm^2$ (**Figure 1C (iii-iv)**). Our results showed that the z-resolution of 42.79±1.12 μm while the curing depth varied between 42.79±1.12 μm to 122.24±4.45 μm for an exposure time range of 0.5 to 2 seconds. Then, several commercially available resins were used to print 3D structures. **(Figure S3). Table 1** provides details about resin formulations and printing conditions while **Figure S4 and S5** provide contact angles (θ1) for all the resins used in this work on PDMS and plasma-etched PDMS. Wettability patterns (cross-shaped) on PDMS window was performed using oxygen plasma etching (Plasma etch Inc.) for 5 minutes at RF power of 75W.

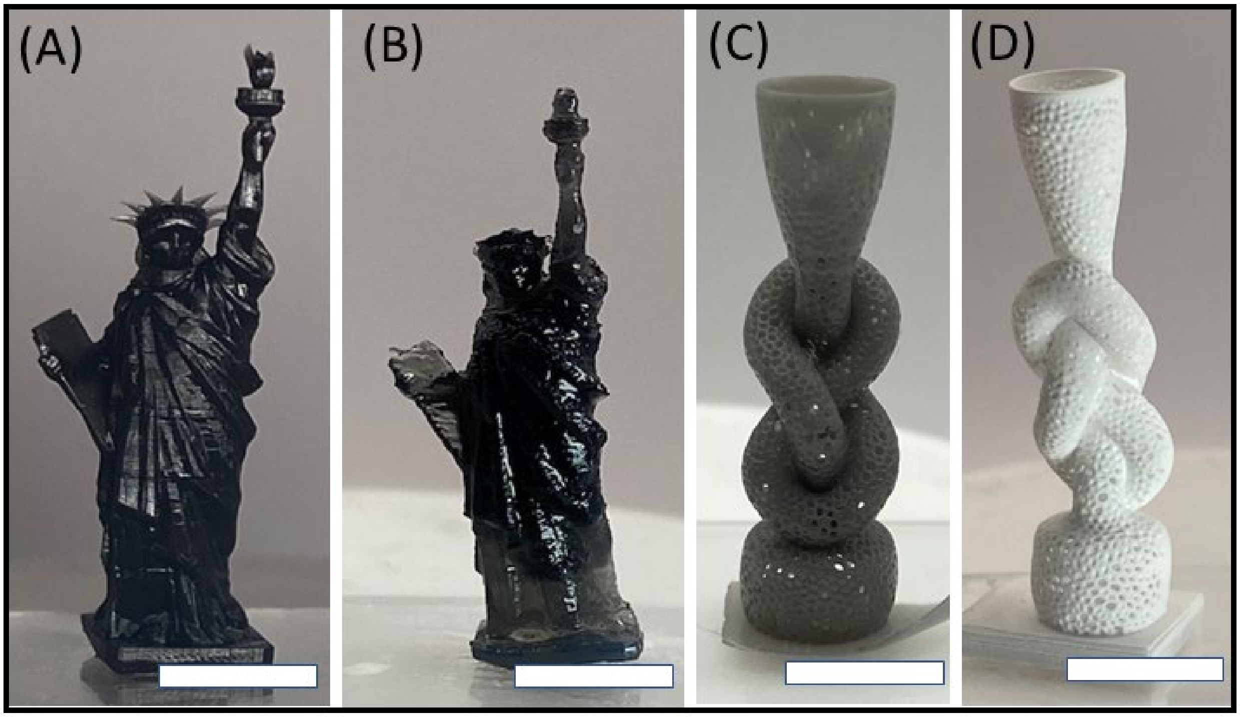


**Figure S3.** MAPS printed 3D structures using different commercially available resins (A) Black resin (B) Flexible-X (C) Photocentric grey (D) Composite-X (Scale bars - 5 mm).

| Photopolymers | Photo-initiator | UV absorber | Intensity (mW/cm$^2$) | Exposure time (seconds) | Viscosity(cps) |
|---|---|---|---|---|---|
| **7.5% GelMA** | 1% LAP | - | 4.4 | 1 | 1.38 |
| **50% PEGDA 700** | 1% LAP | 0.1% tartrazine | 4.4 | 1.3 | 8.16 |
| **PEGDA 6k** | 1% LAP | 0.1% tartrazine | 4.4 | 2.8 | 3.98 |
| **PEGDA 250** | 0.25% Irgacure | 0.5% ITX | 4.4 | 0.8 | 25 |
| **PEGDA 400** | 0.25% Irgacure | 0.5% ITX | 3.25 | 0.7 | 57 |
| **Black resin B9** | - | - | 4.4 | 2.5 | 300 |
| **Flexible - X** | - | - | 4.4 | 3 | 950 |
| **Composite - X** | - | - | 4.4 | 3 | 1400 |
| **Grey resin** | - | - | 4.4 | 2 | 230 |

Table 1. Resin formulation, optimized laser intensity, exposure time and viscosity

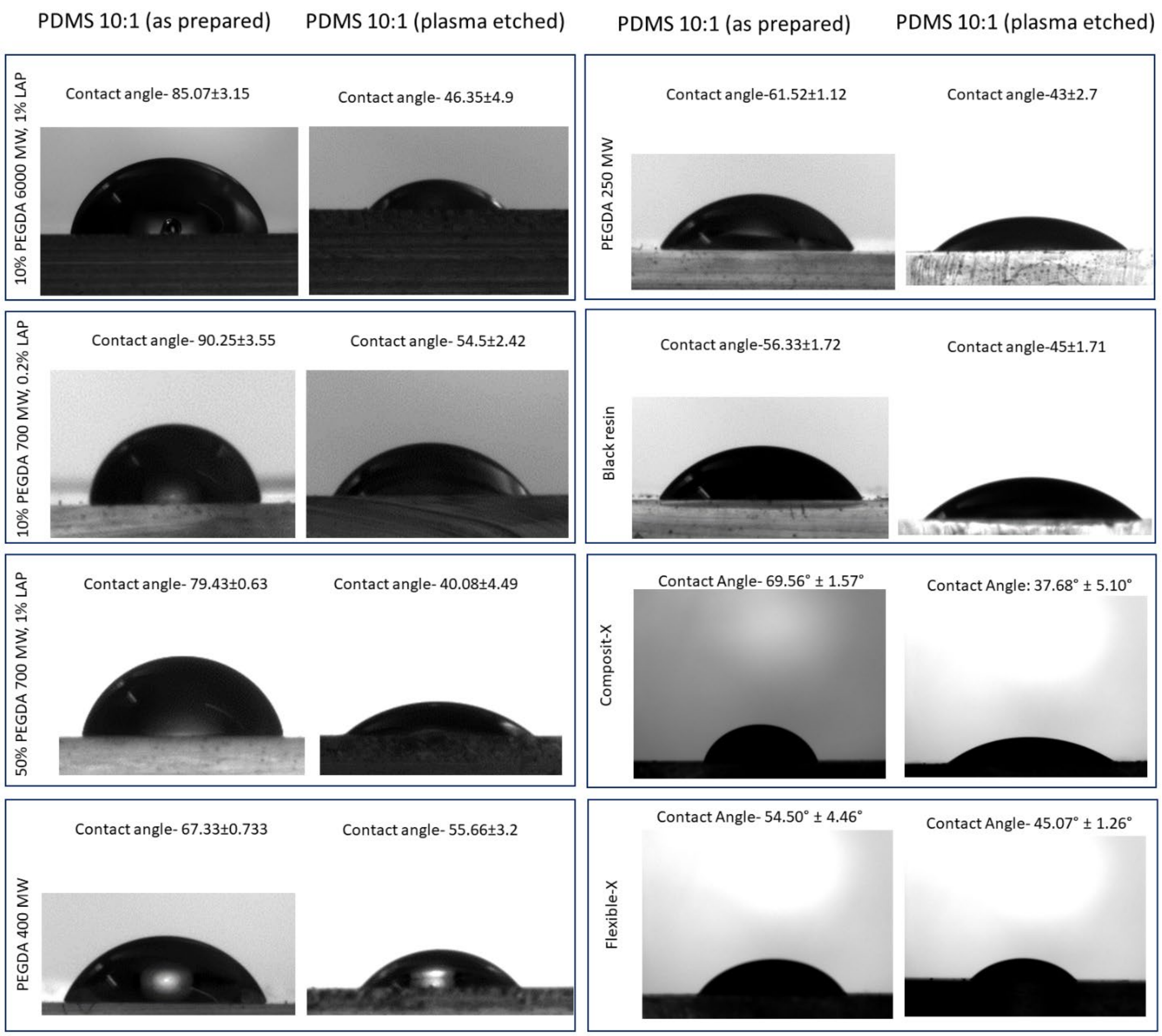


**Figure S4.** Contact angle measured for different photopolymers on native PDMS and oxygen plasma-treated PDMS.

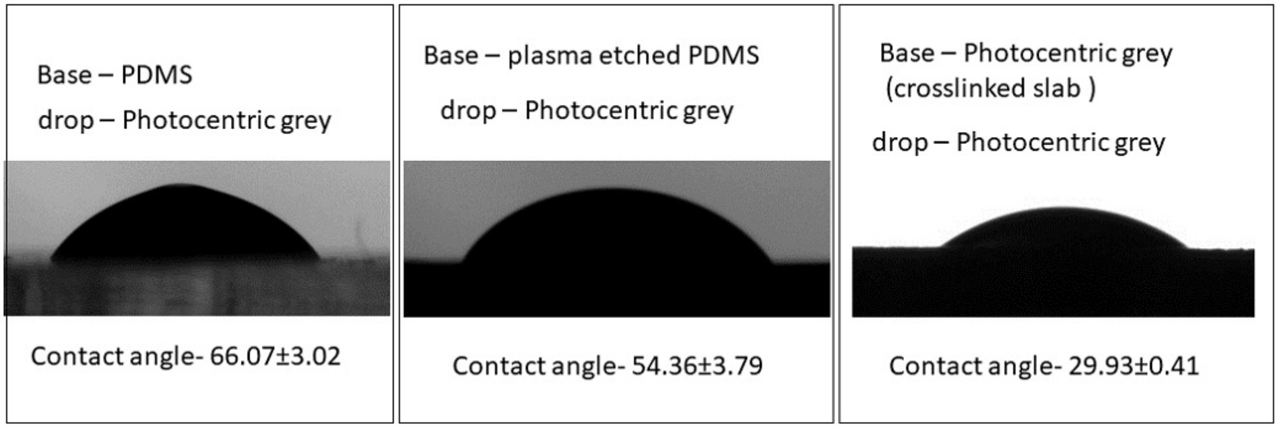


**Figure S5**. Contact angle (θ1) measured for photo-centric grey photopolymer on native PDMS and oxygen plasma-treated PDMS. The contact angle (θ2) for this polymer was also measured on the photo crosslinked slab of the polymer. Further, the contact angle (θ2) was also measured for PEGDA photopolymers, which showed complete wetting of crosslinked slab.

### S5. Many-Body Dissipative Particle Dynamics to simulate the MAPS process

Many-body dissipative particle dynamics (mDPD) method is developed based on standard DPD framework. The time evolution of a mDPD particle $i$ with unit mass $m_i \equiv 1$ is governed by conservation of momentum, which is described by the following set of equations,

$$\frac{d\boldsymbol{r}_i}{dt} = \boldsymbol{v}_i \quad (1)$$

$$\frac{d\boldsymbol{v}_i}{dt} = \boldsymbol{F}_i = \sum_{j \neq i}\left(\boldsymbol{F}_{ij}^C + \boldsymbol{F}_{ij}^D + \boldsymbol{F}_{ij}^R\right) \quad (2)$$

where $t$, $\boldsymbol{r}_i$, $\boldsymbol{v}_i$and $\boldsymbol{F}_i$ denote time, and position, velocity, force vectors, respectively. The summation of forces is carried out over all other particles within a cutoff radius $r_c$, beyond which the direct interactions between particles are considered to be zero.

The three components of $\boldsymbol{F}_i$ include the conservative force $\boldsymbol{F}_{ij}^C$, dissipative force $\boldsymbol{F}_{ij}^D$ and random force $\boldsymbol{F}_{ij}^R$, which are expressed as

$$\boldsymbol{F}_{ij}^C = A\omega_c(r_{ij})\boldsymbol{e}_{ij} + B(\rho_i + \rho_j)\omega_d(r_{ij})\boldsymbol{e}_{ij} \quad (3)$$

$$\boldsymbol{F}_{ij}^D = -\gamma\omega_D(r_{ij})(\boldsymbol{e}_{ij} \cdot \boldsymbol{v}_{ij})\boldsymbol{e}_{ij} \quad (4)$$

$$\boldsymbol{F}_{ij}^R = \delta\omega_R(r_{ij})\xi_{ij}\Delta t^{-\frac{1}{2}}\boldsymbol{e}_{ij} \quad (5)$$

where $r_{ij}$ is distance between particles $i$ and $j$, $\boldsymbol{e}_{ij}$ is the unit vector from particle $j$ to $i$, and $\boldsymbol{v}_{ij} = \boldsymbol{v}_i - \boldsymbol{v}_j$ is the velocity difference. $\omega_c$, $\omega_d$, $\omega_D$ and $\omega_R$ are the weight functions of $\boldsymbol{F}_{ij}^C$, $\boldsymbol{F}_{ij}^D$and $\boldsymbol{F}_{ij}^R$, respectively. A negative coefficient $A < 0$ stands for an attractive force and a positive coefficient

$B > 0$ results in a density-dependent repulsive force. $\gamma$ is dissipative parameter, $\xi$ is the Gaussian white noise with zero mean and unit variance which describes the degrees of freedom that have been eliminated from the coarse-graining process. The dissipative force and random force act as a thermostat if the dissipation parameter $\gamma$ and the amplitudes of white noise $\delta$ satisfy the fluctuation-dissipation theorem requiring $\delta^2 = 2\gamma k_B T$ and $\omega_D(r) = [\omega_R(r)]^2$, in which $k_B$ is the Boltzmann constant and $T$ is the temperature. A common choice of the weight function is $\omega_c = 1 - r_{ij}/r_c$ and $\omega_D = {\omega_R}^2 = \left(1 - r_{ij}/r_c\right)^s$ for $r_{ij} \le r_c$ and vanish for $r_{ij} > r_c$, while $\omega_d = 1 - r_{ij}/r_d$ with a different cutoff radius $r_d < r_c$.

The mDPD system a MAPS model is constructed with parameters $A_{ss} = A_{ll} = -40, B = 25, \gamma = 18, \delta = 6, k_B T = 1, r_c = 1.0, r_d = 0.75, s = 0.5$, and $A_{sl}$ changes from $-26$ to $-34$ to adjust the wettability of solid surfaces, in which the parameters $A_{ss}$, $A_{ll}$ and $A_{sl}$ represent the attractive coefficients in Eq. (3) for solid-solid, liquid-liquid and solid-liquid interactions. Varying the strength of attractive interactions between solid and liquid particles can tune the solid-liquid interfacial tension and hence change the wetting contact angles. The mDPD system with an initial meniscus volume of 11.88 microliters consists of 793,746 particles, in which 169,169 particles form the cross-linked printhead, 198,888 particles being the PDMS substrate, and 425,689 particles representing the liquid meniscus. The parameters in the mDPD model are calibrated based on the fluid properties of plasma etched PDMS (10:1) with a viscosity of $17.41\ \mathrm{mPa \cdot s}$ and a mass density of $960\ \mathrm{kg/m^3}$, and the wetting contact angle $\theta_1$ varies from 40° to 90° for different PDMS base while $\theta_2$ is 29° for crosslinked slab.

**S6. Design and fabrication of microfluidic mixer**

To print gradient structures, a microfluidic mixer, designed and printed using MAPS, was used to precisely mix user-defined resin formulations before infusing droplets into the fabrication window. **(Figure S6).** The micromixer was designed using SOLIDWORKS with two inlets, one outlet, and internal up-and-down features to facilitate mixing of multiple resin solutions (**Figure S6B(i,ii)**). A custom-written MATLAB code was used to slice the micromixer CAD design and converted it to a computer-generated digital masks **(Figure S6B(iii))** which were modulated by DMD to generate light patterns and print the mixed using PEGDA 250 resin. **(Figure S6B(iv))** The rapid mixing process was recorded by flowing two colored PEGDA resin solution from two inlets. (**Figure S7**

**and Video V5)** By adjusting the relative flow rates of the resins, a desired concentration profile can be programmed into the fluidic pumps. **Figure S7(i-iii)** illustrates an example of the flow and mixing of PEGDA 700 MW resin (green) and PEGDA 700 MW (milky-white) resin, where the flow rate for milky-white resin linearly decreased from 0.3 ml/min to 0 ml/min over a period of 18 minutes, while the flow rate of the green resin was reversed. Intensity of the green resin, recorded at the outlet, showed a linear increase over time, indicating efficient mixing. **(Figure S7(iv)).**

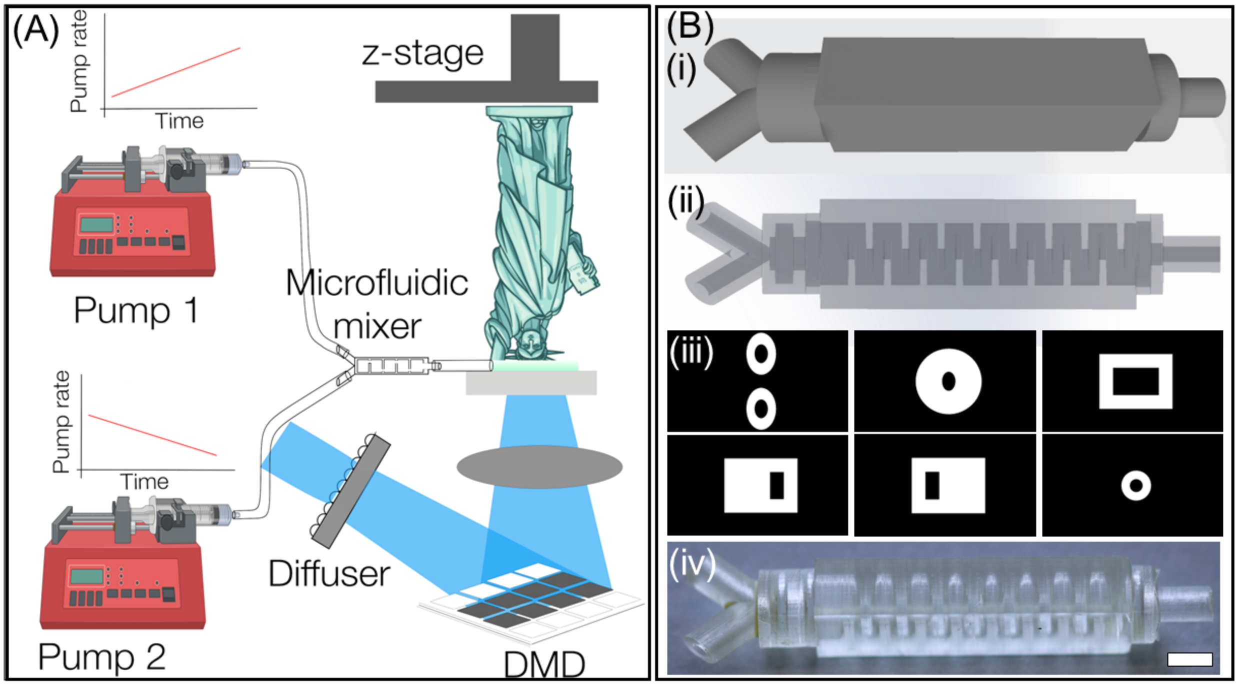


**Figure S6.** (A) Configuration of MAPS during gradient printing. B (i-ii) CAD of the microfluidic mixer showing both overall structure and internal mixing geometry. (iii) Computer-generated masks and as-printed microfluidic mixers using MAPS and PEGDA 250 resin. (iv) (Scale bar-2mm).

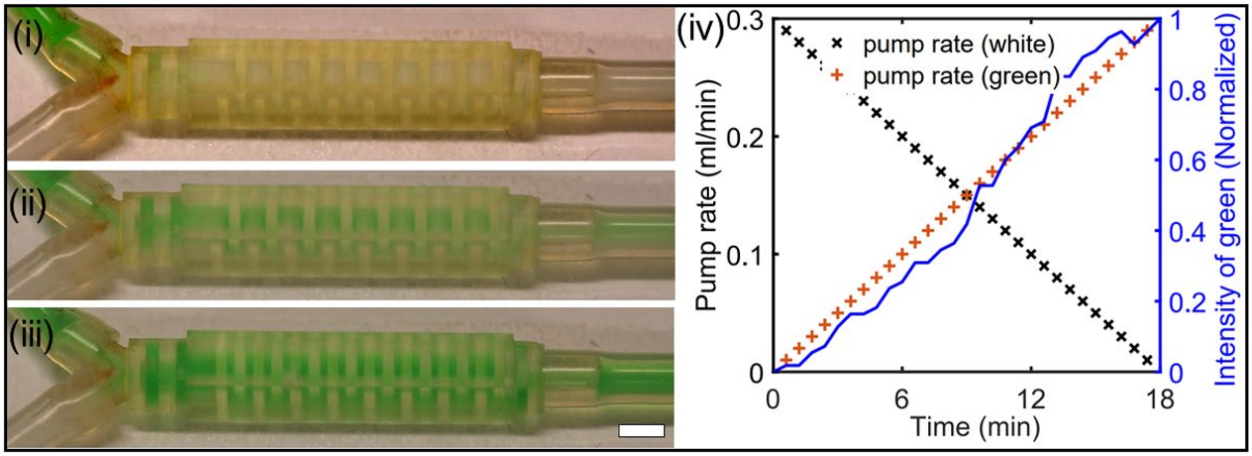


**Figure S7.** The performance of the microfluidic mixer is evaluated using two PEGDA 700 resins (green and milky-white). Recorded images at different time-points are shown: (i) t=0.5 min, (ii) t=9 min, and (iii) t=18 min (Scale bar- 2mm). (iv) A plot illustrates the flow rates of milky white and green resins, while blue line, which records the intensity of green dye at the outlet channel, indicates the mixing efficiency.

### S7. Optical and mechanical characterization of MAPS printed structures

MAPS printed structures were analyzed and imaged using a digital optical microscope (HIROX, KH-8700). For imaging, the MXG-2016Z/MX-2016Z 20-160x objective lens was utilized, providing a resolution of 1.35µm. Fluorescence images were obtained using Leica DMI4000B inverted microscope. Images and Videos were also recorded using a DSLR camera (Cannon EOS T6i). The compression test was conducted at room temperature using an Instron Universal Testing System (Model-5966). Sections of gradient structures were sliced and compressed at a rate of 0.5 mm/min.

### S8. Synthesis of $Fe_3O_4$ nanoparticles and Carbon black

**Chemical**. Iron(II) chloride tetrahydrate ($FeCl_2 \cdot 4H_2O$, 98%), iron(III) chloride hexahydrate ($FeCl_3 \cdot 6H_2O$, 97%), poly(acrylic acid sodium salt) (PAA, average MW ~5100, powder), and ammonium hydroxide solution ($NH_4OH$, 30-33%) were purchased from Sigma Aldrich and used as received without further purifications. Carbon black (VULCAN XC72R, 50 nm, 99%) was purchased from Fisher and used as received without further purifications.

**Synthesis of $Fe_3O_4$ nanoparticles.** The modified co-precipitate method from the previous report is applied to the synthesis of hydrophilic $Fe_3O_4$ nanoparticles (NPs).[4] 6.28 mmole of $FeCl_2 \cdot 4H_2O$, 11.40 mmole of $FeCl_3 \cdot 6H_2O$, and 170 ml of deionized (DI) water were mixed and stirred in a flask. After the $FeCl_2 \cdot 4H_2O$ and $FeCl_3 \cdot 6H_2O$ were fully dissolved, the clear brown solution was heated to 363K on a hot plate. While heating, 9.24 ml $NH_4OH$ and 15.40 ml $PAA_{(aq)}$ were prepared aside for the injection. $NH_4OH$ was used as received. 1.86 g of PAA and 15.4 ml DI water were mixed for the $PAA_{(aq).}$ The solutions were added to the reaction flask via a syringe pump at the rate of 10 ml/min. When the system reached the reaction temperature, $NH_4OH$ was first added to the flask via the syringe pump. Black slurry resulted in the reaction flask after all $NH_4OH$ was added to the flask. The reaction solution was annealed at 363K for 10 mins and then $PAA_{(aq)}$ was added for the surface modification. The reaction was left for another 60 minutes at 363K to complete the PAA coating. After that, the reaction was cooled down for further purification. $Fe_3O_4$ NPs were collected by centrifuging the crude solution at 5000 rpm for 10 mins. The supernatant was discarded, and the precipitate, $Fe_3O_4$ NPs were re-dispersed in DI water. The processes were repeated twice to remove the free PAA and any reaction leftovers. Purified $Fe_3O_4$ NPs were dispersed in DI water.

**S9. Cell culture and staining**

The SOAS-2 cell line is a type of osteosarcoma cell line. To maintain the cells in culture, a medium containing DMEM (Dulbecco's Modified Eagle's Medium) with high glucose and pyruvate was used. This medium was supplemented with 10% fetal bovine serum (FBS), 1% L-glutamine, and 1% penicillin-streptomycin, which provided the necessary nutrients and growth factors to support cell growth and survival.

Specifically, MC3T3 cells were cultured in a 175 $cm^2$ flask until they reached 80% confluency, after which they were harvested. The cells were detached from the flask using trypsin and centrifuged to form a pellet. Subsequently, the pellet was resuspended in cell growth media to achieve a cell concentration of approximately 1 million cells per ml. The final cell suspension was loaded into a sterile syringe to prepare for the printing process.

The MC3T3 cells were cultured in aMEM (Minimum Essential medium) with ribonucleosides, deoxyribonucleosides, 2 mM L-glutamine and 1 mM sodium pyruvate, but without ascorbic acid supplemented 10% FBS and 1% penicillin-streptomycin.

For osteoblastic differentiation, the same cell media was supplemented with 100um Ascorbate (C, L-ascorbic-2-phosphate, Sigma #A8690) and 2mM B-glycerophosphate (BGP, Sigma G9422).

#### *Fluorescence staining*

To perform fluorescence staining on a bioprinted structure, the structure was washed multiple times in PBS before being fixed with 4% formaldehyde for 20 minutes at room temperature. Following fixation, the structure was washed again with PBS and then permeabilized with 0.2% Triton X-100 for 15 minutes at room temperature. The structure was then immunostained with 1:1000 dilution of DAPI for nucleus and 1:400 Alexa Fluor 488 Phalloidin for F-actin in PBS containing 5% horse serum. The immunostained sample was washed and mounted on with EverBrite™ Mounting Medium on glass bottom petri dish and sealed with parafilm. The immunostained sample was imaged using an inverted fluorescence microscope with a DAPI and GFP filter sets.

#### *Alizarin red staining*

The fixed sample was submerged in Alizarin Red Solution (Sigma-Aldrich 2003999) for 2 minutes and washed with PBS several times until the PBS solution was clear. The sample was imaged using a histology microscope.
